# VERY LONG-CHAIN ACYL-CoA SYNTHETASE-3 (ACSVL3) PROMOTES THE MALIGNANT GROWTH BEHAVIOR OF U87 GLIOMA CELLS VIA CHANGES IN CELL CYCLE WITHOUT AFFECTING APOPTOSIS

**DOI:** 10.1101/2023.05.04.539403

**Authors:** Haiyan Yang, Xiaohai Shi, Elizabeth A. Kolar, Emily M. Clay, Shuli Xia, Zhengtong Pei, Paul A. Watkins

## Abstract

Decreasing the expression of very long-chain acyl-CoA synthetase 3 (ACSVL3) in U87MG glioblastoma cells by either RNA interference or genomic knockout (KO) significantly decreased their growth rate in culture, as well as their ability to form rapidly growing tumors in mice. U87-KO cells grew at a 9-fold slower rate than U87MG cells. When injected subcutaneously in nude mice, the tumor initiation frequency of U87-KO cells was 70% of that of U87MG cells, and the average growth rate of tumors that did form was decreased by 9-fold. Two hypotheses to explain the decreased growth rate of KO cells were investigated. Lack of ACSVL3 could reduce cell growth either by increasing apoptosis, or via effects on the cell cycle. We examined intrinsic, extrinsic, and caspase-independent apoptosis pathways; none were affected by lack of ACSVL3. However, significant differences in the cell cycle were seen in KO cells, suggesting arrest in S-phase. Levels of cyclin-dependent kinases 1, 2, and 4 were elevated in U87-KO cells, as were regulatory proteins p21 and p53 that promote cell cycle arrest. In contrast, lack of ACSVL3 reduced the level of the inhibitory regulatory protein p27. γ-H2AX, a marker of DNA double strand breaks, was elevated in U87-KO cells, while pH3, a mitotic index marker, was reduced. Previously reported alterations in sphingolipid metabolism in ACSVL3-depleted U87 cells may explain the effect of KO on cell cycle. These studies reinforce the notion that ACSVL3 is a promising therapeutic target in glioblastoma.

## INTRODUCTION

Gliomas are the largest category of primary central nervous system (CNS) tumors, accounting for 78% of all primary intrinsic malignant CNS tumors (1). Glioblastoma multiforme (GBM), the highest grade of primary brain tumor (World Health Organization grade IV), is the most common and fatal type. GBM tumors grow quickly and are difficult to treat due to the cell type heterogeneity (2). Prognosis is very poor once diagnosed. Even with standard-of-care treatment, most of these patients experience tumor recurrence (3), and none of the current treatments can effectively prolong survival after relapse (4). Therefore, new treatment strategies and new therapeutic targets are highly desirable.

Fatty acids (FA) are fundamental building blocks of phospholipids and sphingolipids. For incorporation into these complex lipids, FA must first be “activated” by thioesterification to coenzyme A (CoA). This reaction is catalyzed by members of a 26-enzyme family of acyl-CoA synthetases (ACS) (5). The ACS family contains subfamilies that have preferences for FA substrates of different acyl chain lengths. We found that ACSVL3, one of 6 members of the very long-chain subfamily, to be overexpressed in gliomas of all grades (6), as well as all types of lung tumor (7). ACSVL3 (gene symbol SLC27A3) is also known as FA transport protein 3 (FATP3).

U87MG cells with knockout of ACSVL3 expression – U87-KO cells – slowed cell proliferation and decreased the number of colonies formed in soft agar. When injected either subcutaneously or orthotopically into nude mice, U87-KO cells produced fewer, slower growing xenografts than did control U87MG cells (6,8). These observations of normalized growth and lack of tumorigenicity of U87-KO cells suggest that ACSVL3 is a promising therapeutic target in GBM and perhaps other human malignancies. Therefore, we sought to understand the mechanism(s) by which depleting U87MG cells of ACSVL3 has beneficial effects on both the in vitro and in vivo growth properties of these cells. Several possibilities were considered, including changes in lipid metabolism, carbohydrate metabolism, autophagy, apoptosis, and cell cycle activity.

We previously reported finding significant alterations in lipid metabolism in U87-KO cells. While no effects on phospholipid synthesis were observed, significant alterations in sphingolipid metabolism were seen when ACSVL3 was depleted in U87MG cells (8). Somewhat surprisingly, substantial changes in carbohydrate metabolism (glycolysis, tricarboxylic acid cycle, and pentose phosphate pathway) were also observed in U87-KO cells (Kolar EA, Shi X, Clay EM, and Watkins PA, unpublished observations). In this report, we address the potential role(s) for apoptosis and cell cycle alteration in the malignant phenotype of U87MG cells. While we find no evidence that ACSVL3 KO in U87 cells promoted apoptosis, significant alterations in the cell cycle, and the expression and phosphorylation of many cell cycle proteins, were observed. The possibility that autophagy alters U87MG cell growth when ACSVL3 is deleted remains under investigation.

## MATERIALS AND METHODS

### Reagents and general methods

The following antibodies were all obtained from Cell Signaling Technology; catalog numbers are in parentheses: caspase 3 (9662), caspase 8 (9746), cleaved caspase 9 (9501), apoptosis-inducing factor (4642), cyclin-dependent kinase (CDK) 1 (9112), CDK2 (2546), CDK4 (2906), cyclin B1 (4135), cyclin D1 (2926), cyclin E1 (4129), p21 (2947), p27 (3686), p53 (9282), checkpoint kinase 2 (CHEK2) (6334), ataxia telangiectasia mutated (ATM) protein kinase (2873), phospho-histone H3 (pH3; Ser10) (9701), and phospho-histone H2A variant X (γ−H2AX) (9718). Horseradish peroxidase-conjugated secondary antibodies for western blot were from Santa Cruz Biotechnology (sc-2357). Alexafluor 488-conjugated secondary antibodies for immunofluorescence were from Jackson ImmunoResearch (211-545-109). Protease inhibitor cocktail (539131) and phosphatase inhibitor cocktail (524625) were both from Calbiochem. Annexin V (A13201) was from Thermo Fisher. Etoposide (GR-307) was from BioMol. Muse® cell analyzer kits for measurement of caspase3/7 (MCH100108) and cell cycle (MCH100106) were from EMD Millipore. Unless otherwise noted, protein was measured by the method of Lowry et al. (9). Statistical significance was determined using Student’s t-test.

### Cell lines and culture

The U87 MG wild type (U87MG) cell line was obtained from American Type Culture Collection (Manassas, VA). U87MG cells were authenticated within a year of their use by the Johns Hopkins Genetics Resources Core Facility by short tandem repeat analysis using the PowerPlex 16 HS kit (Promega). Unless otherwise indicated, all cell culture components were from Corning Cellgro. Cells were cultured in MEM (minimum essential medium, Eagle) containing 10% fetal bovine serum (FBS; Gemini Bioproducts) and supplemented with 10 mM HEPES, 1 mM pyruvate, and nonessential amino acids. Cells were incubated at 37°C in a humidified incubator with a 5% CO_2_ atmosphere. The U87MG ACSVL3 knock out (U87-KO) cell line was produced using zinc finger nuclease technology as previously described (8).

### Western blot analysis

Cells grown in 10 cm culture dishes were washed with phosphate-buffered saline (PBS) and scraped in 100 μl of lysis buffer (25 mM Tris, pH 7.6, 150 mM NaCl, 1% Nonidet-P40, 1% Na deoxycholate, and 0.1% sodium dodecyl sulfate) containing protease inhibitors and phosphatase inhibitor. Cell lysates were prepared in gel loading buffer (10) such that 20 μl contained 30 μg cell protein. Samples were boiled for 5 min and subjected to sodium dodecyl sulfate polyacrylamide gel electrophoresis (SDS-PAGE) on either 10% gels (cast freshly) or 4-20% gradient gels (Invitrogen). Proteins were transferred to polyvinylidene fluoride (PVDF) membranes using an Invitrogen semi-dry blotting apparatus. Membrane proteins were stained with 0.1% Ponceau-S and photographed to verify evenness of transfer; only those membranes where protein bands from U87 and ACSVL3 KO cells were equally intense were processed further. Membranes were blocked by incubation either 1 h at room temperature or overnight at 4oC in Tris-buffered saline + Tween (19 mM Tris, pH 7.4, 137 mM NaCl, 2.7 mM KCl, and 0.1% Tween-20; TBST) containing 10% nonfat dry milk. Incubation with primary antibody (generally 1:1000 dilution in TBST+10% milk) was overnight at 4°C. After washing three times with TBST, membranes were incubated with the appropriate species-specific horseradish peroxidase–conjugated secondary antibody. Signal detection was achieved with SuperSignal West Pico (Thermo Fisher) and Kodak X-Omat AR film. Multiple exposures of each blot were obtained and densitometry of subsaturating exposures was done using ImageJ Fiji (11). For each primary antibody, the density of the band from U87 cell was set to 1.0 and the relative intensity from ACSVL3 KO cells calculated. Data from 3 or more biological replicates were plotted as mean ± standard deviation. Statistical significance was calculated using Student’s t-test.

### Caspase 3/7 activation

Analysis of non-synchronized cells was done using the Muse® cell analyzer and the Caspase-3/7 Kit. This allows for rapid, quantitative measurements of apoptotic status based on Caspase-3/7 activation, and cellular plasma membrane permeabilization and cell death. The manufacturer’s instructions were followed to obtain the relative percentage of cells that are live, apoptotic, and dead.

### Immunofluorescence analysis and annexin V staining

Cells were cultured under normal conditions on glass coverslips and grown to 40-60% confluence. For immunofluorescence, cells were fixed with 3% formaldehyde, permeabilized with 1% Triton X-100, and incubated with primary antibodies and fluorescent-conjugated secondary antibodies (see Materials) as previously described (12). The mounting medium contained DAPI to stain nuclei. Images were captured using a Zeiss Axio Imager M2 Fluorescence Microscope with Apotome using identical exposure conditions. ImageJ (imagej.net) was used to quantitate mitotic index (pH3) and DNA damage (γH2AX). Annexin V staining was performed using the manufacturer’s instructions (Thermo Fisher), and visualization was as described for immunofluorescence.

### Proteomic analysis

Proteomic profiling was performed exactly as previously described (8,13,14). Data from biological replicates was obtained and the fold change in U87-KO vs. U87MG was calculated.

### Cell cycle

Analysis of non-synchronized cells was done using the Muse® cell analyzer. U87MG and U87-KO cells were harvested from 10 cm culture dishes using trypsin. Cells were counted using a hemocytometer. A total of ∼1 × 10^6^ cells were used for each sample. Samples were labeled and processed as described by the Muse Cell Cycle Assay kit (EMD Millipore). Briefly, cells were washed and resuspended in a final volume of 50 μl with PBS. The suspension was added dropwise into 1 ml ice cold 70% ethanol while vertexing. Ethanol-fixed cells were washed with PBS, resuspended in Muse Cell Cycle Reagent and incubated for 30 min in the dark before analysis. Gating was adjusted after the first sample was analyzed and maintained throughout the run. The instrument was set to acquire 10,000 events per sample. Data were analyzed using Muse® software.

Cells were synchronized by incubation in serum-free culture medium for 48 hrs. The block was released by switching to culture medium containing 10% FBS. Cells were harvested for cell cycle analysis at 0, 6, 12, 18, and 24 hours post FBS addition as previously described (19).

## RESULTS

### U87-KO cells do not show evidence of increased apoptosis by annexin V staining

There are several possible mechanisms to explain why depleting U87MG cells of ACSVL3 has beneficial effects on growth and tumorigenesis, including effects on membrane lipid synthesis, apoptosis, autophagy, and cell cycle. In this work, we investigated two of these – apoptosis and cell cycle. To assess whether the decrease in malignant growth properties of ACSVL3-depleted U87 cells could be due to increased apoptotic cell death, we examined several indices of apoptosis.

Phosphatidylserine (PS), a phospholipid normally found in the inner leaflet of the plasma membrane, migrates to the outer leaflet in apoptotic cells. Fluorescein isothiocyanate (FITC) tagged annexin V binds to PS on the surface of apoptotic cells. When cells were incubated under normal culture conditions (10% FBS) and stained with this early apoptotic marker, there was no evidence of apoptosis in either U87MG (Fig. 1, top panel, A) or U87-KO (Fig. 1, top panel, B) cell lines. To verify that the assay was functioning normally, we treated cells overnight with reduced (0.1%) FBS plus 40 μM etoposide prior to annexin V staining. As shown in Fig. 1, top panel, C and D, this treatment induced apoptotic changes in a small percentage of cells, indicating that both parental and KO U87MG cells are relatively resistant to apoptosis.

**Figure 1.**
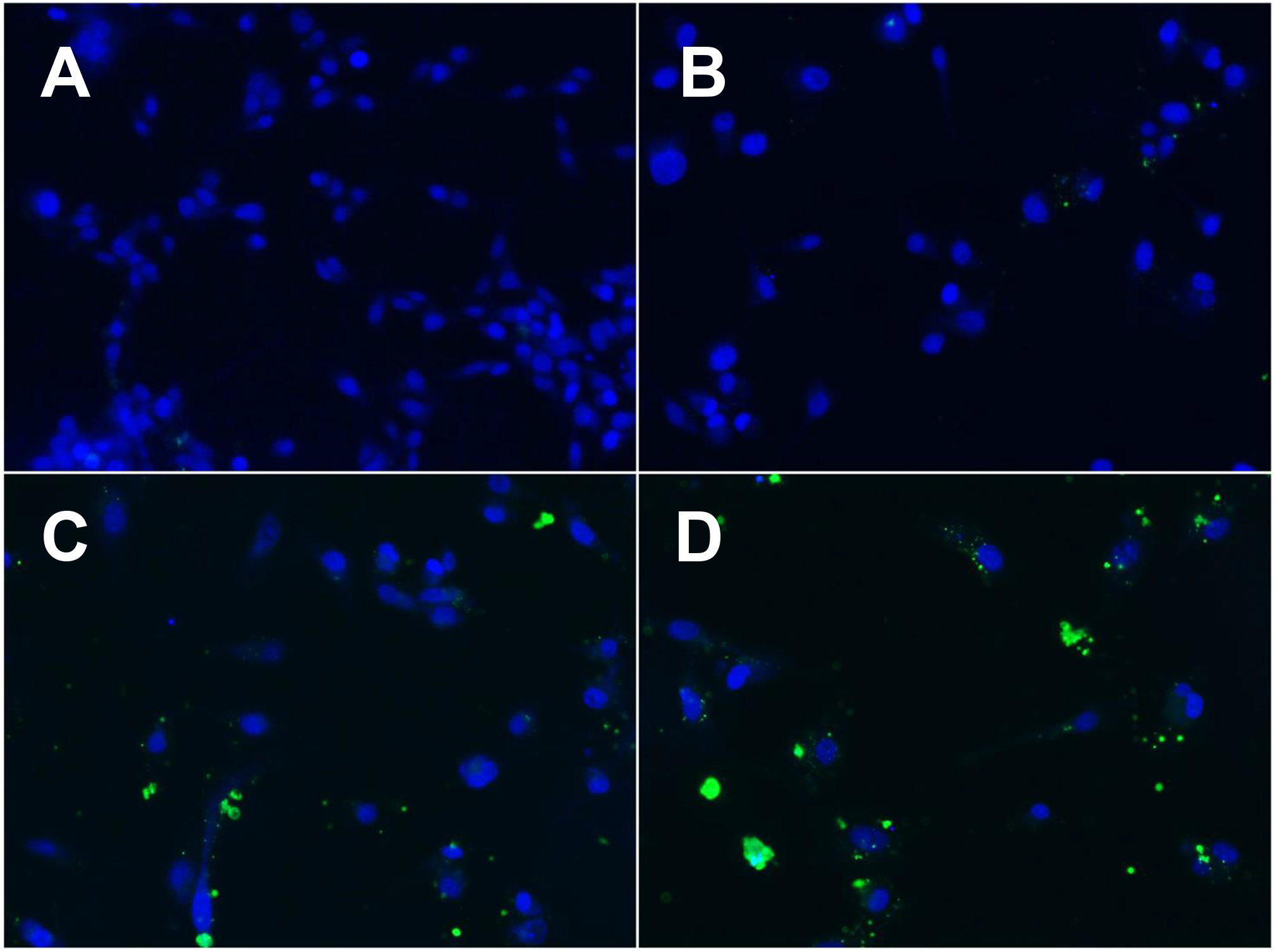
KO of ACSVL3 does not increase annexin V staining. U87MG (A) and U87-KO cells (B) were cultured under normal conditions and then incubated with annexin V as described in Methods. Cell nuclei were stained with DAPI (blue dye). Under normal growth conditions (A and B), less than 1% of cells showed evidence of annexin V binding (green dye). As a positive control, U87MG (C) and U878-KO cells (D) were cultured overnight in low (0.1%) FBS culture medium containing 40 μM etoposide to induce apoptosis. Green fluorescence was observed in several cells, indicating that the staining method was functioning properly.

### KO of ACSVL3 does not promote caspase activation by either extrinsic or intrinsic apoptotic pathways

We next determined if KO of ACSVL3 induced activation of caspase 8 (extrinsic apoptosis pathway), caspase 9 (intrinsic pathway), or caspase 3 (extrinsic and intrinsic). Western blotting showed no evidence of the 18 kDa active caspase 8 cleavage product in either U87MG (Fig. 2A, lane 1) or U87-KO cells (Fig. 2A, lane 2) incubated under normal culture conditions. A band corresponding to the 43 kDa cleavage product was barely detectable in U87-KO cells. Overnight treatment of U87MG cells with 40 μM etoposide plus 40 ng/ml TNF_α_ yielded a small amount of both 18 kDa and 43 kDa caspase 8 fragments (Fig. 2A, lane 3), demonstrating that the antibody would have detected cleavage products had they been present. The negligible amount of caspase 8 cleavage indicates that the low growth rate of U87-KO cells is not due to increased apoptosis via the extrinsic pathway. Under normal culture conditions, no 37 kDa active caspase 9 cleavage product was detected in either U87MG cells (Fig. 2B, lane 1) or U87-KO cells (Fig. 2B, lane 2). When U87MG cells were treated overnight with 40 μM etoposide plus 40 ng/ml TNF_α_, a band corresponding to the active, cleaved caspase 9 was seen (Fig. 2B, lane 3), again demonstrating that the antibody would have detected an increase in this marker of intrinsic apoptosis had it been the cause of the decreased growth rate of ACSVL3-deficient cells. Finally, neither U87MG cells (Fig. 2C, lane 1) nor U78-KO cells (Fig. 2C, lane 2) showed evidence of a 17 kDa active caspase 3 fragment when cultured under normal conditions. A small amount of cleaved product was detected after overnight incubation of U87MG cells with 40 μM etoposide plus 40 ng/ml TNF_α_ (Fig. 2C, lane 3), again indicating that the antibody would have detected this product had apoptosis been induced by ACSVL3 KO.

**Figure 2.**
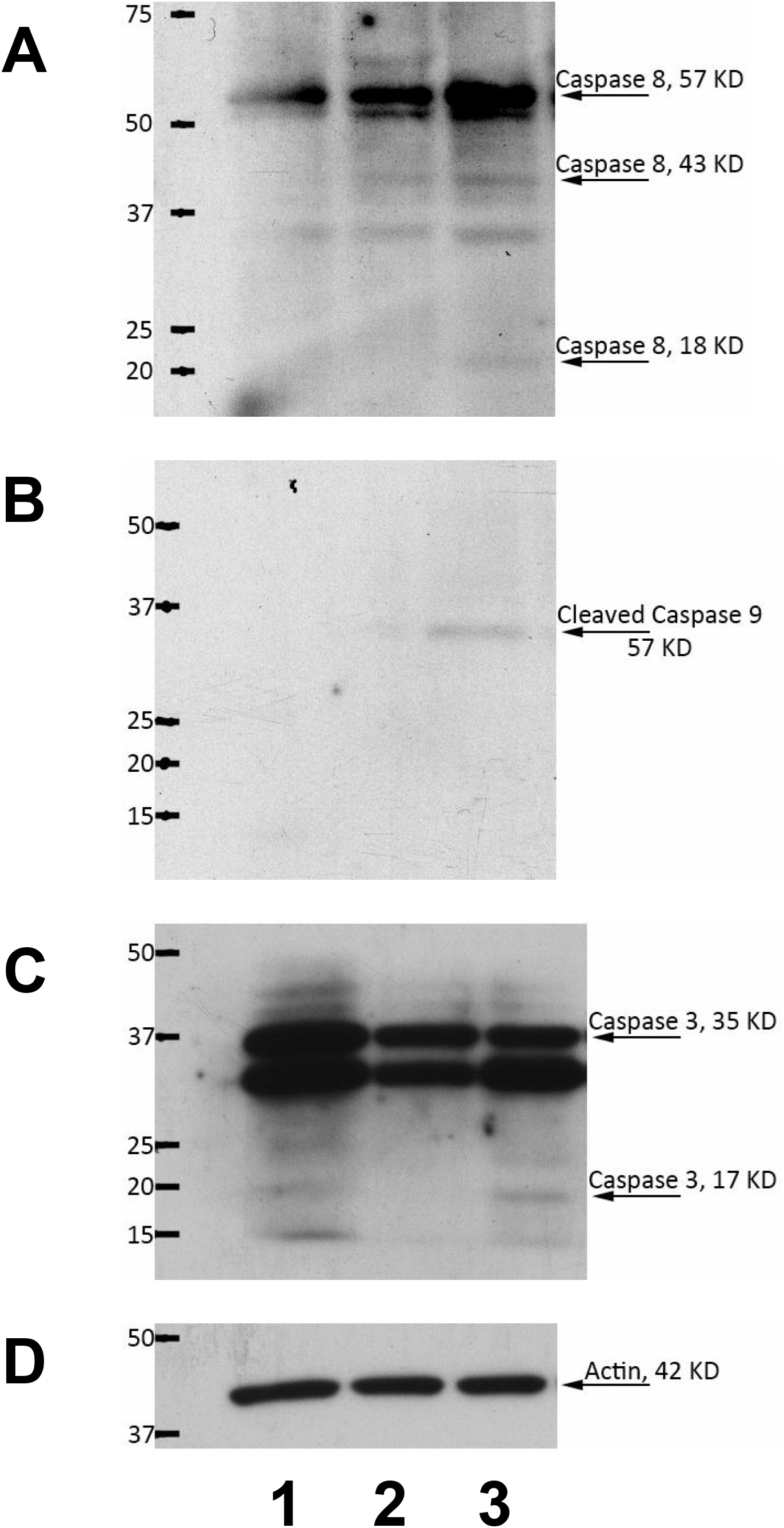
Caspases 8, 9, and 3 are not activated in ACSVL3 KO cells. Cells were scraped in lysis buffer and Western blotting was performed as described in Methods. To generate positive controls, U87 cells were incubated overnight in reduced-serum (0.1% FBS) medium containing etoposide (40 μM) and TNFα (40 μM). For all blots, lane 1 contains U87MG cells, lane 2 contains U87-KO cells, and lane 3 contains etoposide- and TNFα-treated U87MG cells. This study was repeated at least 3 times; a representative blot is shown. A, caspase 8 cleavage (apoptosis extrinsic pathway) yields an 18 kDa fragment seen only in lane 3. B, caspase 9 (apoptosis intrinsic pathway) cleavage yields a 57 kDa fragment only seen in lane 3. C, caspase 3 cleavage yields a 17 kDa fragment primarily seen in lane 3. D, actin loading control. There was no evidence for increased apoptosis (lane 2 fragments) in U87-KO cells for any of the three caspases studied.

### Neither caspase 3/7 activation nor non-caspase-dependent apoptosis are affected by ACSVL3 KO

Using the Muse® Cell Analyzer, flow cytometry was used as an independent measure of caspase 3/7 activation; this assay also quantitates the number of live vs. dead cells. As shown in Fig. 3, top panel, 98-99% of cells were alive and non-apoptotic for both U87MG and U87-KO cells. The number of live apoptotic U87MG cells was not statistically different from U87-KO cells and was < 1.5% of total live cells. The percent of dead cells never exceeded 1%. These data are in agreement with the lack of caspase activation determined by Western blot in Fig 2. Apoptosis inducing factor (AIF) is a mitochondrial protein involved in non-caspase-dependent cell death (16). During induction of apoptosis, AIF is released from the intermembrane space and translocates to the nucleus, where it induces chromatin condensation and DNA fragmentation. Immunofluorescence analysis of U87MG and U87-KO cells using an AIF antibody (Fig. 3, bottom panel, A and B) showed no nuclear staining, and no differences between the two cell types. Incubation of cells with 0.1% FBS plus 40 μM etoposide prior to AIF immunostaining showed an increased intensity of mitochondrial fluorescence, but again without nuclear staining (Fig. 3, bottom panel, C and D). These observations suggest that caspase-independent apoptosis is not increased in ACSVL3-deficient cells.

**Figure 3.**
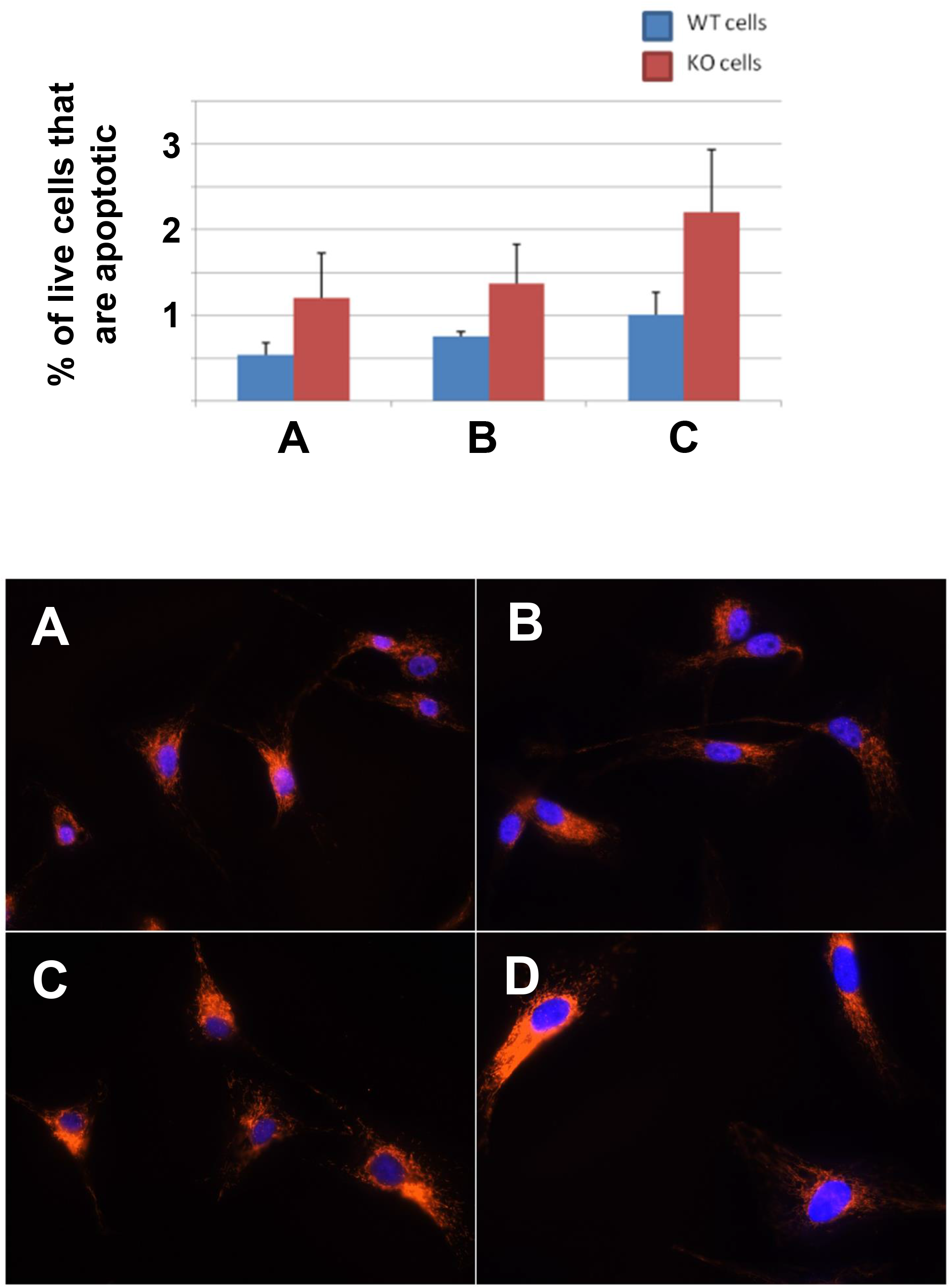
Caspase 3/7 activation and mitochondrial release of AIF are not increased in ACSVL3 KO cells. Top panel. Flow cytometry with a Muse® cell analyzer was used to separate live, apoptotic live, dead, and apoptotic dead cells based on caspase 3/7 activation. Results presented are the average of 3 independent analyses ± SD. (A) Under normal growth conditions, less than 2% of total U87MG and U87-KO cells were either dead or dead and apoptotic. >98% of both U87MG and U87-KO cells were live and non-apoptotic. There was a small but insignificant increase in the % of live apoptotic cells in ACSVL3-deficient U87 cells vs. U87MG cells (1.2% vs. 0.5%). In an attempt to induce apoptosis, cells were incubated in medium containing either reduced FBS (0.1%) (B), or reduced FBS + 40 μM etoposide (C). While reduced FBS alone had no effect on apoptosis, the combination of reduced FBS plus etoposide marginally increased the percent of live apoptotic cells. Bottom panel. Apoptosis inducing factor (AIF) translocated from mitochondria to the nucleus in apoptotic cells. Immunofluorescence using an antibody to AIF showed no difference between U87MG cells (A) and U87-KO cells (B). Treatment with reduced serum (0.1% FBS) plus etoposide increased mitochondrial staining but did not promote nuclear translocation in either U87MG cells (C) and ACSVL3 KO cells (D). Both studies reinforce the conclusion that U87MG cells are highly resistant to apoptosis, and that KO of ACSVL3 in these cells did not render them more apoptotic.

### The cell cycle is altered by ACSVL3 depletion in U87MG cells

Another potential cause of growth rate reduction in ACSVL3 KO U87MG cells and xenografts is changes in cell cycle. To assess this possibility, we used flow cytometry of both non-synchronized and synchronized cells. U87MG cells maintained under normal culture conditions (i.e., non-synchronized cells) showed were found to be mainly in G0/G1 phase. In contrast, U87-KO cells were mainly in S phase (Fig. 4A). To pursue this further, cells were synchronized by serum starvation for 48 hours, after which they were returned to complete medium containing 10% FBS. At the onset of serum restoration, both U87MG and U87-KO cells had a similar cell cycle profile, with the majority of cells in G0/G1 phase (Fig. 4B). By 12 hours post-serum restoration, however, significantly more U87-KO cells were found in S phase and fewer in G0/G1, while the percent of U87MG cells in G0/G1 remained almost unchanged.

**Figure 4.**
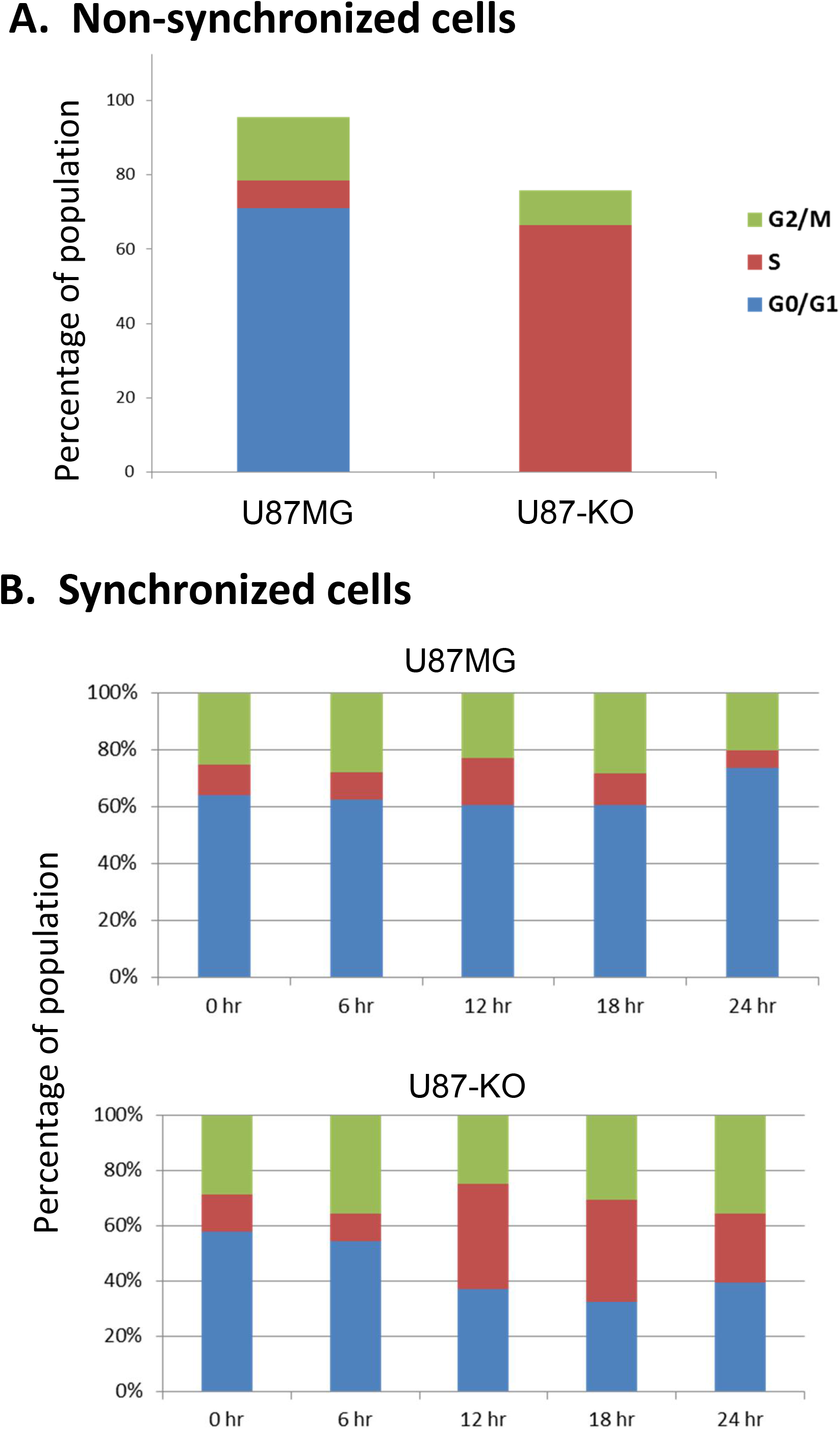
KO of ACSVL3 in U87MG cells significantly alters the cell cycle. Top panel. Cells grown to near confluence under normal culture conditions (non-synchronized) were harvested and the cell cycle assessed using a Muse® cell analyzer as described in Methods. The majority of U87MG cells were in G0/G1, while most U87-KO cells were in S-phase. The cell cycles of U87MG cells and ACSVL3-deficient cells (lower panel) were synchronized by a 48 hr of serum starvation. Cells were either harvested (time 0 hr) or returned to medium containing 10% FBS and then harvested after incubation for 6, 12, 18, and 24 hr. Cell cycle analysis of propidium iodide-stained cells was performed as previously described (19). While the cell cycle profiles of both U87MG and U87-KO cells were similar at 0 hr, divergence of KO cells toward the pattern seen in unsynchronized cells was evident by 12 hr following serum re-supplementation.

### Cell cycle protein levels were altered in U87-KO cells

To gain further understanding of these differences, we assessed the levels of several cell cycle proteins. Cyclin-dependent kinases (CDK) 1, 2, and 4 were all elevated in ACSVL3 KO cells (Fig. 5); the increases were statistically significant (CDK1, p<0.01; CDK2 and 4, p<0.05). Cyclins B1, D1, and E1, which are regulatory subunits of CDK1, 4 and 2, respectively, were also elevated (Fig. 5); these increases were also statistically significant (all p<0.05). The cell cycle is exquisitely regulated by many proteins, including p53, p21/CDKN1A (CDK inhibitor 1A), and p27/CDKN1B (CDK inhibitor 1B).

**Figure 5.**
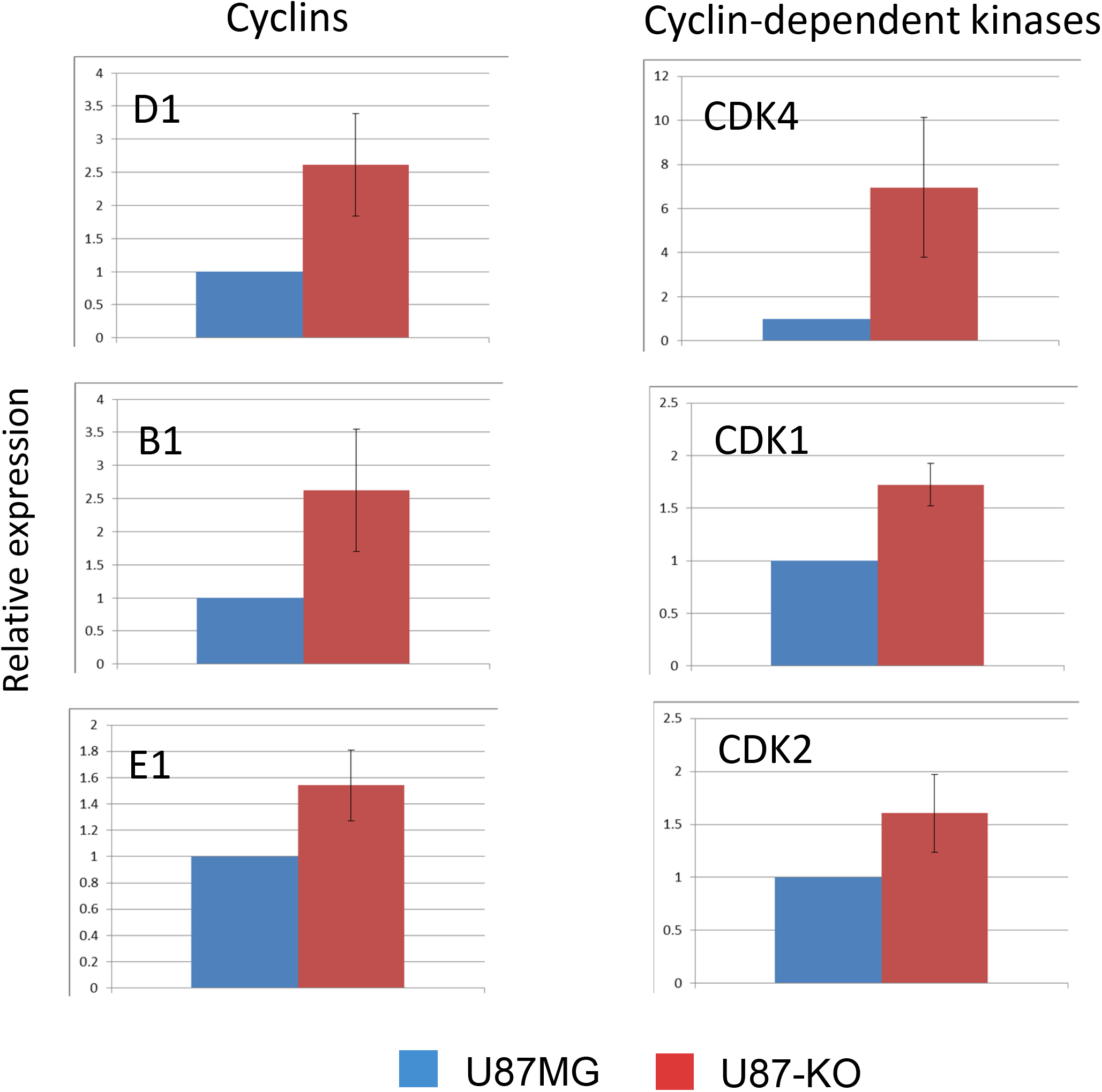
Cyclins and cyclin-dependent kinases (CDKs) are upregulated in ACSVL3 KO cells. Western blotting was used to determine expression levels of several CDKs and their respective regulatory partner cyclins. Non-synchronized cells were grown to near confluence before harvesting. Results presented are the average of 3 independent analyses ± SD. Expression levels of cyclin D1 and CDK4, cyclin B1 and CDK1, and cyclin E1 and CDK2 were all elevated in U87-KO cells relative to U87MG cells. All changes were statistically significant (p<0.05). level.

While levels of p53 and p21, both of which promote cell cycle arrest, were higher in KO cells than in U87MG cells, levels of inhibitory p27 were lower (Fig. 6); all results were statistically significant (p<0.05). Proteomic analysis detected several cyclins, CDKs, and cell cycle regulatory proteins in U87MG and ACSVL3 KO cells (Table 1). Although several proteins of interest were not sufficiently abundant for detection by mass spectrometry, e.g. p21 and p27, the levels of most of cell cycle-related proteins were higher in KO cells than in U87MG cells. Thus, the proteomics data confirm the western blot data.

**TABLE 1.** Expression of cell cycle proteins in U87-KO cells vs. U87MG cells.

| **Gene** | **Protein** | **Fold-change**<br>**U87-KO/U87MG** |
| --- | --- | --- |
| CDK1 | Cyclin-dependent kinase 1 | 2.2 |
| CDK2 | Cyclin-dependent kinase 2 | 2.0 |
| CDK2AP2 | CDK2-associated protein 2 | 2.4 |
| CINP | CDK2 interacting protein | 1.3 |
| CDK5 | Cyclin-dependent kinase 5 | 2.3 |
| CDK5RAP1 | CDK5 regulatory subunit-associated protein 1 | 0.8 |
| CDKAL1 | CDK5 regulatory subunit-associated protein 1-like 1 | 1.3 |
| CDK5RAP2 | CDK5 regulatory subunit-associated protein 2 | 0.8 |
| CDK5RAP3 | CDK5 regulatory subunit-associated protein 3 | 3.0 |
| CDK11B | Cyclin-dependent kinase 11B | 1.5 |
| CDK12 | Cyclin-dependent kinase 12 | 2.0 |
| CDK13 | Cyclin-dependent kinase 13 | 3.5 |
| CDKN2AIP | Cyclin-dependent kinase inhibitor 2A-interacting protein | 3.9 |
| CCNH | Cyclin H | 5.1 |
| CCNK | Cyclin K | 4.0 |
| CCNT1 | Cyclin T1 | 5.9 |
| CCNY | Cyclin Y | 0.5 |
| CCNYL1 | Cyclin Y-like protein 1 | 0.8 |
| CKS1B | Cyclin-dependent kinases regulatory subunit 1 | 2.6 |
| CKS2 | Cyclin-dependent kinases regulatory subunit 2 | 1.7 |
| CCNB1 | G2/mitotic-specific cyclin B1 | 1.9 |
| CCNB2 | G2/mitotic-specific cyclin B2 | 1.0 |

Proteomic analysis detected several cyclins, cyclin-dependent kinases, and other cell cycle-associated proteins. Expression of most of these proteins was higher in ACSVL3 KO cells relative to U87MG cells.

**Figure 6.**
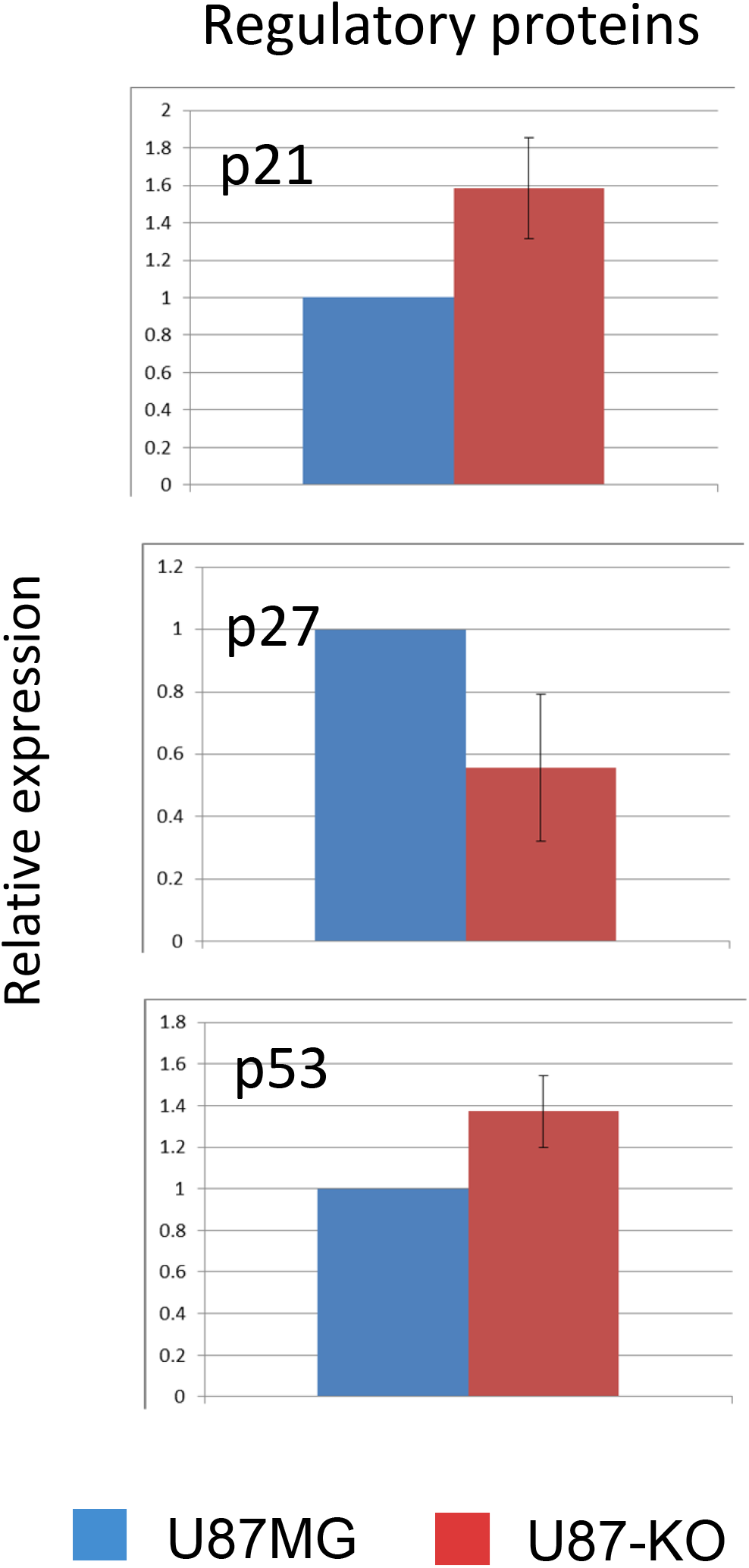
Expression levels of cell cycle regulatory proteins are altered by KO of ACSVL3. Western blotting was used to determine expression levels of several other proteins that regulate the cell cycle as described in the legend to Figure 5. While p21 and p53 expression was higher in ACSVL3-deficient cells, expression of p27 was lower. All changes were statistically significant (p<0.05).

### Depletion of ACSVL3 in U87MG cells reduces DNA double-strand breaks and lowers the mitotic index

These alterations in cell cycle proteins could be the result of DNA damage in KO cells. Phosphorylated H2A histone family member X (γ-H2AX) is a biomarker for double-strand DNA breaks. As shown in Fig. 7A, there was increased γ-H2AX staining in cells lacking ACSVL3. The mitotic index of cells can be assessed by immunostaining with antibody against phosphorylated histone H3. In agreement with cell growth curves and cell cycle data, the mitotic index of U87-KO cells was significantly lower than in U87MG cells (Fig. 7B).

**Figure 7.**
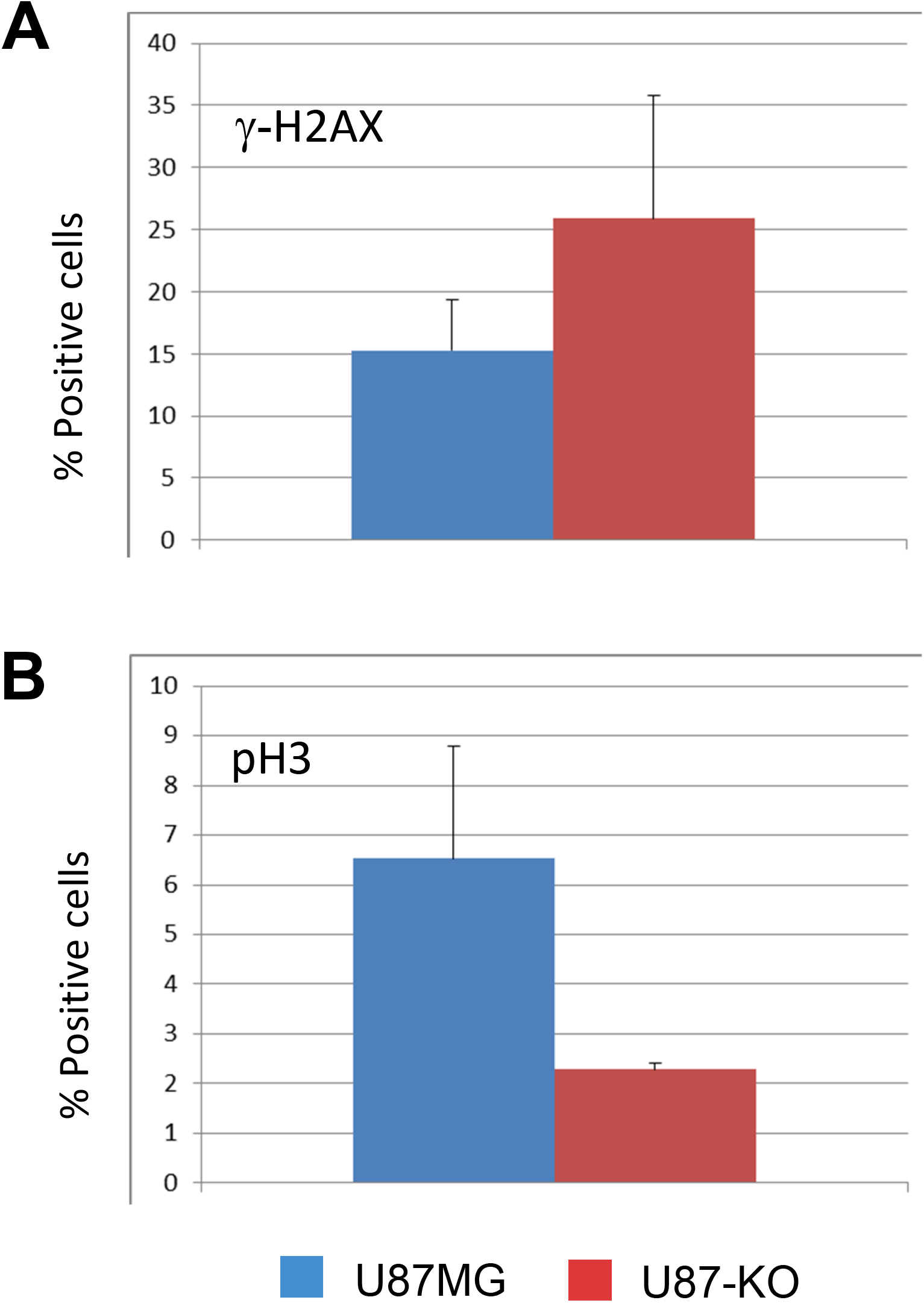
Markers of mitotic index and DNA damage markers are altered in ACSVL3 KO cells. Immunofluorescence was used to assess evidence for DNA double-strand breaks and changes in mitotic index in U87MG cells and U87-KO cells. Top. Phosphorylation of histone H2A family member X (γ-H2AX) is a biomarker for double-strand DNA breaks. Immunostaining with antibody specific for γ-H2AX revealed a higher percentage of positive cells when ACSVL3 was depleted. Bottom. Phosphorylation of histone H3 (pH3) is a marker of mitotic index. Immunostaining with pH3-specific antibody revealed a lower mitotic index in ACSVL3-depleted U87 cells.

## DISCUSSION

Previous reports from our laboratory suggested that ACSVL3 – an enzyme whose sole known function is to esterify coenzyme A to a fatty acid substrate – is a promising therapeutic target in both glioblastoma and lung cancer (6-8). When ACSVL3 was knocked down by RNA interference in glioblastoma cells or lung cancer cells, they grew slower in monolayer culture and produced fewer colonies in suspension culture. Glioblastoma cells in which ACSVL3 was knocked down produced fewer, slower-growing tumors when implanted in mice either subcutaneously or orthotopically. These beneficial effects of knockdown (6,7) were recapitulated in U87-KO cells in which the ACSVL3 gene was knocked out using ZFN technology (8).

Our overall goal is to characterize U87-KO cells to gain further insight into glioma malignancy. There are several potential mechanistic explanations for the beneficial effects of ACSVL3 depletion, including metabolic changes, alterations in signaling pathways, autophagy, cell cycle, increased apoptosis, or combinations of these possibilities. In this work, we focused on apoptosis and the cell cycle.

We first asked whether knockout of ACSVL3 rendered U87MG cells more apoptotic. Apoptosis, or programmed cell death, is a highly regulated process that becomes dysregulated in pathological conditions such as cancers (reviewed in (17)). Control of cell number in normal tissues by apoptosis prevents either under- or overgrowth, thus maintaining homeostasis. Decreased apoptosis is a feature of many malignancies. Therefore, it was logical to ask whether depletion of ACSVL3 reduced the malignant properties of U87 cell by increasing the number of apoptotic cells.

We found no evidence that U87-KO cells were more apoptotic that U87MG cells. There was no evidence for translocation of phosphatidylserine from the inner to outer leaflet of the plasma membrane, as evidenced by lack of annexin V staining (Fig. 1). Activation of the intrinsic (mitochondrial) apoptosis pathway was not detected. Neither activation of Caspase 9, a marker of the intrinsic pathway, nor caspase 8, a marker of the extrinsic (death receptor) pathway, was observed (Fig. 2). Caspase 3 is a downstream common effector caspase for both the extrinsic and intrinsic pathways, but its activation was also not detected (Figs. 2 and 3). The assay for caspases 3/7 also measured cell viability, and indicated that >98% of cells were alive for both U87MG and U87-KO (Fig 3). Even after incubation with reduced serum plus etoposide, >97% of U87-KO cells were alive, indicating a significant resistance to apoptosis in the U87 cell line. Finally, there was no evidence of increased translocation of AIF, a marker of caspase-independent apoptosis (18), to nuclei in U87-KO cells. Thus, by all criteria tested, we found no evidence that growth reduction in U87-KO cells results from an increase in apoptosis.

In contrast to the studies on apoptosis, we found rather significant differences in cell cycle parameters between U87MG and U87-KO cells. When cells were grown in a non-synchronous fashion – i.e. normal growth conditions – about 70% of U87MG cells were in G0/G1-phase. This is similar to findings previously reported by other laboratories (Fig. 4) (19,20). In contrast, most of the U87-KO cells were in S-phase. This suggested that major changes in cell cycle could provide an explanation for the growth rate differences between the malignant U87MG cells and the less-malignant U87-KO cells. Synchronizing the cell cycle by serum deprivation added further insight into cell cycle alterations when ACSVL3 is depleted in U87 cells. Just before restoring 10% FBS to the culture medium, U87-KO cells and U87MG cells exhibited a similar distribution between G0/G1, S, and G2/M (Fig. 4, time=0 hrs). However, by 12 hours following serum restoration, many more U87-KO cells were persisting in S-phase, as observed when cells were maintained under normal growth conditions.

The distinction between cell cycle in U87MG vs. ACSVL3-depleted U87-KO cells was further confirmed by changes in expression of several cyclins and their associated cyclin-dependent kinases (CDKs). We measured cyclins and CDKs that regulate entry into early G1 (cyclin D1 and CDK 4), S (cyclin E1 and CDK2), and M (cyclin B1 and CDK1). Somewhat surprisingly, protein levels of all were elevated when ACSVL3 was depleted. However, cyclin levels must be regulated for normal progression to occur. Cell cycle arrest can occur if cyclins are not degraded properly (https://www.nature.com/scitable/topicpage/cdk-14046166/). Protein levels of most cyclins and CDKs detected by unbiased proteomics were elevated by ACSVL3 KO (Table 1). Several proteins of interest, however, were below the level of detection, including cyclins D1 and E1, CDK4, checkpoint kinases CHEK1 and CHEK2, WEE1, ATM/ATR, p21, and p27. Failure of a carefully orchestrated pathway of cyclin and CDK degradation may contribute to the slower, more normal, growth rate in U87-KO cells.

The mechanism by which ACSVL3’s acyl-CoA synthetase enzymatic activity regulates cell cycle has not been definitively identified. Acyl-CoA synthetases in the “very long-chain” ACSVL family (gene symbols SLC27A1-A6) generally activate long-to very long-chain fatty acids (21). We subsequently reported that ACSVL3 (SLC27A3) has a preference for fatty acids containing 18-22 carbon atoms (8). We also found that rather than decrease the supply fatty acids for bulk membrane phospholipids in rapidly dividing U87MG cells, lack of this enzyme in U87-KO cells primarily affected sphingolipid metabolism (8). Levels of ceramides containing 18-22 carbon fatty acids were decreased. Levels of signaling molecules sphingosine and sphingosine-1-phosplate were reduced in U87-KO cells. Although ceramides generally promote cell cycle arrest and apoptosis, its metabolite ceramide 1-phosphate stimulates cell proliferation and invasiveness, and inhibits apoptosis (22). Sphingosine-1-phosphate also promotes proliferation and survival, and can also affect transformation, angiogenesis, cell migration (23,24). Excessive S-1-P is associated with cancer and other human disease pathologies (24). Thus, effects of ACSVL3 depletion on sphingolipid metabolism may in turn affect, in a beneficial way, the cell cycle in U87 GBM cells.

In summary, the lack of ACSVL3 in U87-KO cells reverts them to a more “normal” phenotype. In particular, U87-KO cells grow more like non-transformed cells, and are less tumorigenic. While this cannot be explained by increased apoptosis, changes in cell cycle components are consistent with the improved growth parameters. Changes in cell cycle may be mediated by the altered sphingolipid metabolism observed in U87-KO cells. These observations reinforce our belief that ACSVL3 is a promising therapeutic target in GBM.

## ACKNOWLEDGEMENTS

The authors thank Drs. Akhilesh Pandey and Raja Sekhar Nirujogi, McKusick-Nathans Institute of Genetic Medicine, Johns Hopkins University School of Medicine for proteomic analysis of U87MG and U87-KO cells. This work was supported by NIH grants NS37355 and HD24061.

## REFERENCES

1. Ostrom, Q. T., Price, M., Neff, C., Cioffi, G., Waite, K.A., Kruchko, K., Barnholtz-Sloan, J.S. (2022) CBTRUS Statistical Report: Primary Brain and Other Central Nervous System Tumors Diagnosed in the United States in 2015-2019. Neuro-oncology 24(Suppl 5):v1–v95. doi: 10.1093/neuonc/noac202.

2. Mineo, J. F., Bordron, A., Baroncini, M., Ramirez, C., Maurage, C. A., Blond, S., and Dam-Hieu, P. (2007) Prognosis factors of survival time in patients with glioblastoma multiforme: a multivariate analysis of 340 patients. Acta neurochirurgica 149, 245–252; discussion 252-243 doi: 10.1007/s00701-006-1092-y.

3. Sathornsumetee, S., Reardon, D. A., Desjardins, A., Quinn, J. A., Vredenburgh, J. J., and Rich, J. N. (2007) Molecularly targeted therapy for malignant glioma. Cancer 110, 13–24. DOI: 10.1002/cncr.22741

4. Weller, M., van den Bent, M., Hopkins, K., Tonn, J. C., Stupp, R., Falini, A., Cohen-Jonathan-Moyal, E., Frappaz, D., Henriksson, R., Balana, C., Chinot, O., Ram, Z., Reifenberger, G., Soffietti, R., Wick, W., and European Association for Neuro-Oncology Task Force on Malignant, G. (2014) EANO guideline for the diagnosis and treatment of anaplastic gliomas and glioblastoma. The Lancet. Oncology 15, e395–403. DOI: 10.1016/S1470-2045(14)70011-7

5. Watkins, P. A., Maiguel, D., Jia, Z., and Pevsner, J. (2007) Evidence for 26 distinct acyl-coenzyme A synthetase genes in the human genome. J Lipid Res 48, 2736–2750. doi: 10.1194/jlr.M700378-JLR200

6. Pei, Z., Sun, P., Huang, P., Lal, B., Laterra, J., and Watkins, P. A. (2009) Acyl-CoA synthetase VL3 knockdown inhibits human glioma cell proliferation and tumorigenicity. Cancer Res 69, 9175–9182. doi: 10.1158/0008-5472.CAN-08-4689.

7. Pei, Z., Fraisl, P., Shi, X., Gabrielson, E., Forss-Petter, S., Berger, J., and Watkins, P. A. (2013) Very long-chain acyl-CoA synthetase 3: Overexpression and growth dependence in lung cancer. PLoS One 8, e69392. doi: 10.1371/journal.pone.0069392

8. Kolar, E.A., Shi, X., Clay, E.M., Moser, A.B., Lal, B., Nirujogi, R.S., Pandey, A., Bandaru, V.V.R., Laterra, J., Pei, Z., Watkins, P.A. (2021) Very long-chain acyl-CoA synthetase 3 mediates onco-sphingolipid metabolism in malignant glioma. Med Res Arch 9, 2433. doi: 10.18103/mra.v9i5.2433.

9. Lowry, O. H., Rosebrough, N. J., Farr, A. L., and Randall, R. J. (1951) Protein measurement with the Folin phenol reagent. J. Biol. Chem. 193, 265–275

10. Laemmli, U. K. (1970) Cleavage of structural proteins during the assembly of the head of bacteriophage T4. Nature 227, 680–685. doi: 10.1038/227680a0

11. Schindelin, J., Arganda-Carreras, I., Frise, E., Kaynig, V., Longair, M., Pietzsch, T., Preibisch, S., Rueden, C., Saalfeld, S., Schmid, B., Tinevez, J. Y., White, D. J., Hartenstein, V., Eliceiri, K., Tomancak, P., and Cardona, A. (2012) Fiji: an open-source platform for biological-image analysis. Nat Methods 9, 676–682. doi: 10.1038/nmeth.2019.

12. Watkins, P. A., Gould, S. J., Smith, M. A., Braiterman, L. T., Wei, H.-M., Kok, F., Moser, A. B., Moser, H. W., and Smith, K. D. (1995) Altered expression of ALDP in X-linked adrenoleukodystrophy. Am. J. Hum. Genet. 57, 292–301

13. Nirujogi, R.S., Wright, J.D., Manda, S.S., et al. (2015) Phosphoproteomic analysis reveals compensatory effects in the piriform cortex of VX nerve agent exposed rats. Proteomics 15, 487–99. doi:10.1002/pmic.201400371

14. Kim, M.S., Pinto, S.M., Getnet, D., et al. (2014) A draft map of the human proteome. Nature. 509(7502), 57581. doi:10.1038/nature13302

15. Xia, S., Li, Y., Rosen, E. M., and Laterra, J. (2007) Ribotoxic stress sensitizes glioblastoma cells to death receptor induced apoptosis: requirements for c-Jun NH2-terminal kinase and Bim. Mol Cancer Res 5, 783–792. doi: 10.1158/1541-7786.MCR-06-0433.

16. Bano, D., and Prehn, J. H. M. (2018) Apoptosis-Inducing Factor (AIF) in Physiology and Disease: The Tale of a Repented Natural Born Killer. EBioMedicine 30, 29–37. doi: 10.1016/j.ebiom.2018.03.016.

17. Rebecca S Y Wong Apoptosis in cancer: from pathogenesis to treatment Review J Exp Clin Cancer Res. 2011 Sep 26;30(1):87. doi: 10.1186/1756-9966-30-87.

18. Cregan, S. P., Dawson, V. L., and Slack, R. S. (2004) Role of AIF in caspase-dependent and caspase-independent cell death. Oncogene 23, 2785–2796. doi: 10.1038/sj.onc.1207517.

19. Chockalingam, S. and Ghosh, S.S. (2013) Amelioration of Cancer Stem Cells in Macrophage Colony Stimulating Factor-Expressing U87MG-Human Glioblastoma upon 5-Fluorouracil Therapy. PLoS ONE 8, e83877. doi: 10.1371/journal.pone.0083877.

20. Ye, L., Wang, C., Yu, G., et al. (2014) Bmi-1 induces radioresistance by suppressing senescence in human U87 glioma cells. Oncol Lett 8, 2601–2606. doi: 10.3892/ol.2014.2606.

21. Watkins, P.A. (2008) Very long-chain acyl-CoA synthetases. J. Biol. Chem. 283, 1773–1777. doi: 10.1074/jbc.R700037200

22. Gomez-Larrauri, A., Presa, N., Dominguez-Herrera, A., Ouro, A., Trueba, M., Gomez-Muñoz, A. (2020) Role of bioactive sphingolipids in physiology and pathology. Essays Biochem 64, 579–589. doi: 10.1042/EBC20190091

23. Pyne, N.J. and Pyne, S. (2020) Recent advances in the role of sphingosine 1-phosphate in cancer. FEBS Lett. 594, 3583–3601. doi: 10.1002/1873-3468.13933.

24. Pyne, N.J. and Pyne, S. (2010) Sphingosine 1-phosphate and cancer. Nat Rev Cancer 10, 489–503. doi: 10.1038/nrc2875.

